# Is vectorial transmission of *Trypanosoma cruzi* an efficient route to support high infection rates in sylvatic hosts?

**DOI:** 10.1101/860189

**Authors:** Juan Manuel Cordovez, Mauricio Santos-Vega, Diana Erazo, Camilo Sanabria, Lina María Rendón, Felipe Guhl

## Abstract

Chagas disease is caused by the parasite *Trypanosoma cruzi* and it is transmitted to humans by the triatomine bug *Rhodnius prolixus*. The main insect vector in the Andean countries presents sylvatic and domestic cycles involving humans, insects and reservoirs (e.g small mammals). It is commonly assumed that vectorial transmission is the main route for parasite spread between hosts. Recent studies have reported high percentages (21-80%) of infected opossums (*Didelphis marsupialis*) in the sylvatic cycle, raising the question of whether such a high proportion of infected could be only maintained by vectorial transmission, a seemingly inefficient pathway. To address this question, we formulated a mathematical model that describes the sylvatic transmission dynamics considering vectors and hosts and parametrized with field data. Our results show that vectorial transmission it is not sufficient to explain such high percentages of infected host-mammals reported in the literature. Here we propose oral transmission as an alternate route of transmission that may increase the number of infected individuals found in field studies.

## Introduction

The transmission of the parasite *Trypanosoma cruzi*, etiological agent of Chagas disease, involves several pathways and results in 6 million infected people in Latin America [1]. Human infections are caused by multiple routes, the main suggested mechanism is vectorial transmission that occurs when triatomine insects feed on host blood (Sylvatic mammals and humand after a short period, the vector defecates releasing large amounts of parasites in the skin close to the wound allowing the parasite to reach the bloodstream [2-4]. Vertical transmission occurs in humans from an infected mother to a child; however, the ability of the parasite to cross the placenta of sylvatic reservoirs has not been fully demonstrated yet. Oral transmission has been proved to cause more aggressive clinical symptoms in humans and to have a high mortality rate (8-35% compared to 5-10% by vectorial transmission) only two weeks post infection [5]. In sylvatic mammals, oral transmission has been reported when mammals feed on *Rhodnius prolixu*s infected with *T. cruzi* were ingested [6]. Recent studies in central Brazil have demonstrated that both vertical and oral transmission are not a rare event in this biological system. In fact, a recent study in the Pantanal Region of Brazil, have demonstrated that both the vertical an oral transmission are likely to occur, depending on the encounter possibilities of the mammals and vectors [7].

In sylvatic mammals, particularly of the family Didelphidae, it has also been suggested that spraying from anal glands could be playing an important role in the transmission of *T. cruzi*. Opossums, mainly of the species *D. marsupialis*, have been proposed not only as a reservoir but also as a *T. cruzi* vector, since the parasite can multiply extracellularly in the anal glands of the animal [8-9]. This variety of transmission mechanisms and their relative importance in human infection compared to reservoir infection, suggests that parasites have different transmission cycles in the environment, sylvatic or domestic, that can be connected or isolated depending on the feeding behaviour of insect vectors.

When insect vectors breed and feed inside the houses, a domestic cycle is occurring, involving human and domestic mammals as reservoirs [4,10]. On the other hand, the sylvatic cycle involves triatomine bugs living in the wild feeding on sylvatic mammals such as opossums and rodents that act as reservoirs. The connection between the two cycles occurs when insects migrate from the sylvatic to the domestic habitat attracted by light sources and domestic species presence combined with the increase in the number of domestic animals like dogs [11-14]. It must be considered that the ecological interactions and encounters between vectors and reservoirs depend on the faunal composition, which is directly related with the landscape structure [15].

The domestic and peri domestic cycle has been the focus of many reports [12, 16-17] and control programs [4, 18-20], probably because is the easiest one to intervene and involves humans directly. However, understanding the sylvatic cycle is crucial because it is the source of infected insects that ultimately invade the houses. Furthermore, control programs in Colombia that used pyrethroid insecticides, that succeeded in countries like Chile and Uruguay in eliminating insects from houses [21], have shown limited effectiveness in Colombia due to a strong re-infestation phenomenon that occurs weeks after the application of the insecticide [22]. In addition, vector prevention and control activities have not been very efficient in endemic areas due to the lack of knowledge about the biological characteristics of the vector populations present in each region, leading to uncertainty about the most appropriate control measures in each transmission scenario [23]. Thus, we investigate in more detail the sylvatic cycle dynamics for the Colombian endemic department of Casanare a territory of high interest involving both Chagas disease transmission cycles and high *T. cruzi* natural infection in *R. prolixus* [24-27].

In rural areas of Colombia, particularly in the Orinoco region where the Casanare department is located, *R. prolixus* is considered the main vector of *T. cruzi* transmission. Its main habitat are palm trees, particularly the species *Attalea butyracea*, in a landscape where houses are scarce and well spread. This palm tree species is widely distributed and establishes a large faunistic reserve, where mammalian reservoirs are typically found in the crowns [23]. In this natural habitat, where triatomines and mammals coexist and interact, the vectorial pathway of parasite transmission between them has been demonstrated by many studies [2]. Few authors had documented the existence of both oral and congenital pathways [28-29]. This raises the difficulty of knowing whether vector transmission is acting as a sole route, or if multiple transmission modes act simultaneously.

Regardless of the transmission route, the question of how severe is the infection in sylvatic reservoirs has appeared repeatedly and several studies have reported the proportion of different infected mammals. This is important because high levels of reservoirs infection suggest higher probabilities for human infection. For example, bats can play an important role in transmission scenarios, because their high mobility allows the parasite to migrate between the sylvatic and domestic habitat. Studies carried out in Casanare in different bat species reported infection indexes of T. cruzi from 6.5% to 51% [26, 30]. Moreover, Didelphis marsupialis has been considered one of the main reservoirs of the parasite and previous studies suggest infection rates ranging from 5 to 90% [31-32]. More recent studies have reported 80% and 89% infected mammals in an area with similar ecological characteristics to the department of Casanare [33-34. Finally, the results from our studies showed an infection rate of 21% in *D. marsupialis*. Hypothetically if vectorial transmission is acting alone, host reservoirs will become infected only if after a blood meal the insect defecates on the skin and then the parasites find their way into the bloodstream, and insects would become infected if they suck blood from an infected mammal. This poses the question of whether vectorial transmission is sufficient to sustain more than 21% of infected hosts in a population, as reported before.

Several mathematical models of Chagas disease have been proposed in the last two decades to study its epidemiology and more recently its ecology. Among the first models we can find general analysis of vector and host dynamics [35], the incorporation of acute and chronic stages [36] or models accounting for congenital transmission [37], spatially explicit models [38], age-structure models [39], and stochastic models [40]. On the ecological side, more recent models have investigated the role of dogs [41] and synantropic animals in human infection [42], and the sylvatic cycle dynamics in different habitats varying in their host communities [43]. However, none of these models has evaluated if the assumed mechanisms of transmission could explain the sylvatic host infection that is observed in the wild.

Here we investigate if vectorial transmission per se is capable to maintain the transmission observed in the data by using a mathematical modeling approach. By exploring the model, we will address if the well-known vectorial transmission is enough to support the high infection rates among sylvatic reservoirs and, if so, to propose entomological control strategies that would be adequate to reduce the risk of infection to humans.

## Methods and Results

We formulate different epidemiological models to recreate the sylvatic transmission cycle and establish the interactions between vectors and reservoirs. In the model, we consider hosts (*H*) and vectors (*V*) to represent the population of *D. marsupialis* and *R. prolixus* respectively, and the model was set to represent the dynamics in the region of Casanare, Colombia composed by large palm plantations. The results are normalized per palm as a study unit. The total number of hosts, *N*_*H*_, is divided into susceptible (*S*_*H*_) and infected class (*I*_*H*_). Similarly, the total population of vectors *N*_*V*_, is divided into susceptible (*S*_*V*_) and infected (*I*_*V*_). Our first model depicted in figure 1A, the vectors become infected at a rate (*β*_*HV*_) that reflects the transmission of the parasite from an infected host to a susceptible vector due to biting. Then the host become infected at a rate (*β*_*VH*, 0_) that reflects the transmission of the parasite from an infected vector to a susceptible host due to biting. Likewise, (*β*_*VH*, 1_) is the transmission rate by ingestion (consumption or predation) of the infected vector by the reservoir. In this way *β*_*HV*_, *β*_*VH*, 0_, and *β*_*VH*, 1_ consider triatomine – host contact rate by the probability of infection from the one to the other. The second model, in figure 1B, considers the same routes of transmission as the previous model and includes a new rate reflecting the possibility of susceptible host acquiring the parasite from an infected host (*β*_*HH*_).

The two models are summarized by the equations:

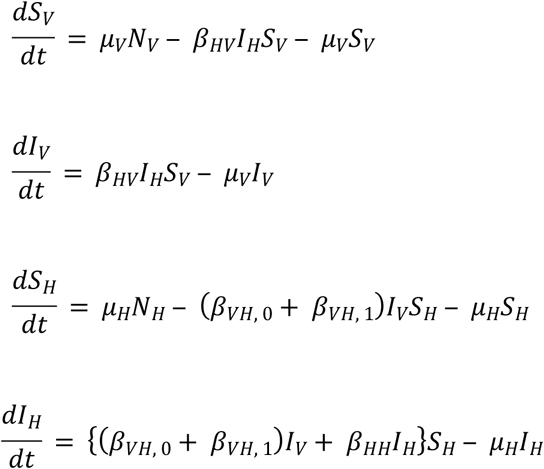

**Figure 1.**
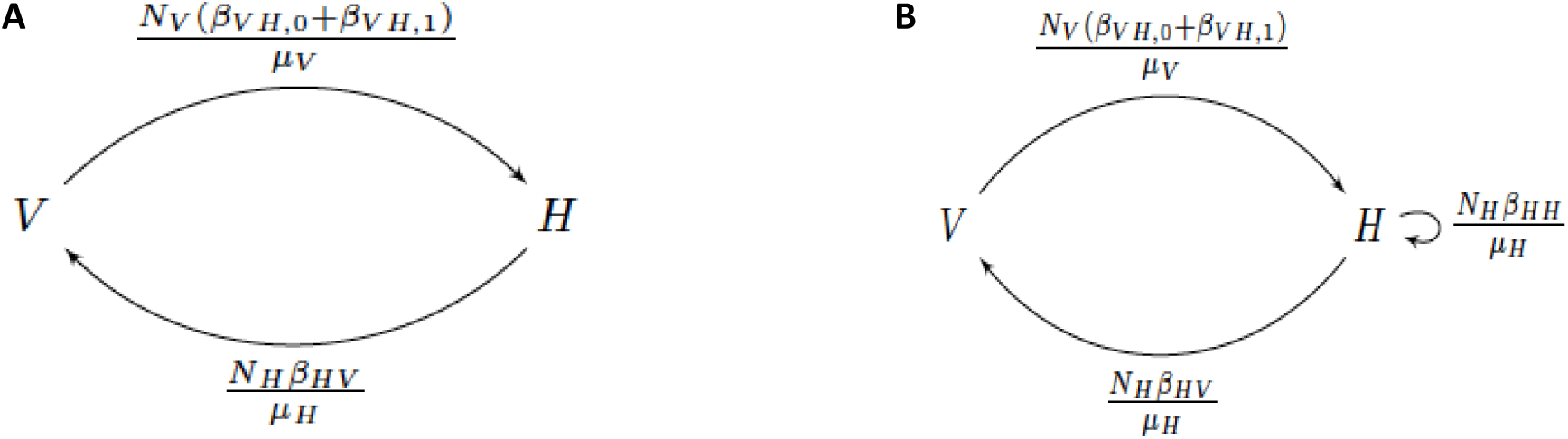
A) Shows the Vector-host model. Vector-host model with transmission between hosts

Where the model with no transmission between hosts (Figure 1A) is obtained by making *β*_*HH*_ = 0. Having no vital dynamics, *N*_*V*_ = *S*_*V*_ + *I*_*V*_ and *N*_*H*_ = *S*_*H*_ + *I*_*H*_ are constant, the equations collapse into:

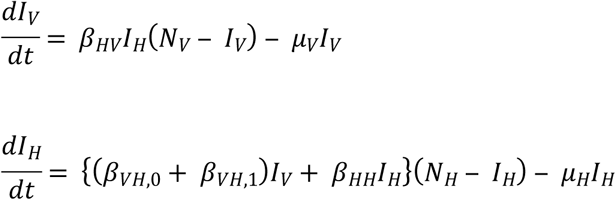

The Next Generation Matrix (*G*) for this model is:

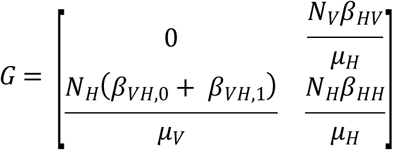

And the adjacency matrix (*S*(*G*)) is then:

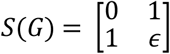

Where *ϵ* = 1 or *ϵ* = 0, depending whether there is transmission between hosts. The spectral radius *ρ* of *S*(*G*) is either 1 if *ϵ* = 0, or 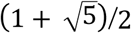 if *ϵ* = 1. Now if *μ*(*G*) is the *critical virulence* [44] of our epidemiological system and *R*_0_ its basic reproductive number:

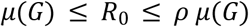

Therefore, with no transmission between hosts

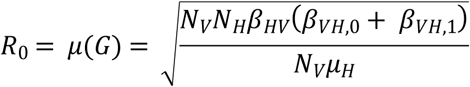

And with it

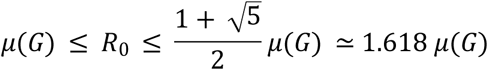

Where

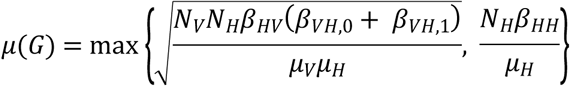

In order to have an endemic equilibrium, where the parasite invaded the ecosystem, it is necessary to have *R*_0_ > 1. In the first model (Figure 1A), in this equilibrium, if it exists, the number of infected hosts corresponds to:

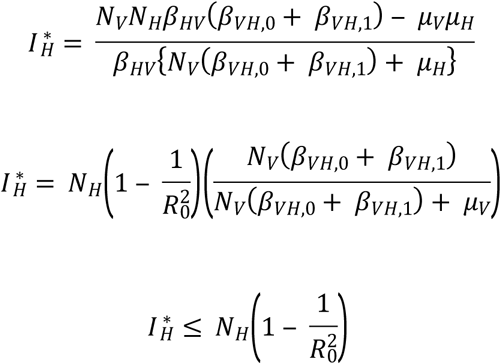

In the second model (Figure 1B), we know that the parasite will not invade the ecosystem if

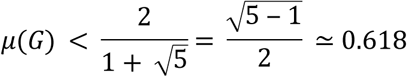

For the numerical exploration of the model we used parameters from the literature. To determine their maximum values for transmission coefficients, which are the most uncertain parameters, we combined expert knowledge, field and lab measures from several sources (Table I). Death rate parameters were estimated based on the average of the life expectancy (1/life expectancy) from literature reports.

Finally, to relax the assumption of the transmission following the mass action law an alternate way to model the transmission would be incorporate saturation effects in the transmission. The most common way to include the effects of saturation in a biological model is to consider an analogous to the Michaelis-Menten equation in the definition of both transmission coefficients. Here, we redefined the transmission rates in our model to include saturation in the following manner:

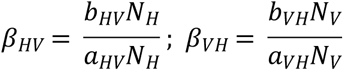

where *b* is the maximum transmission coefficient and *a* reflects the number of hosts necessary to reach the saturation level (when *N*_*H*_ = *a*, the transmission coefficient takes the value of *b*/2). However, even if we consider a more realistic interaction between reservoirs and vectors the model could predicts reproductive numbers higher than 1 for some combinations of the transmission coefficients and does not explain the high levels of infection found in the field studies.

A sensitivity analysis for global reproductive number R0 was performed using the Latin Hypercube method (LH) to estimate each parameter contribution. Negative Partial Rank Correlation Coefficients (PRCC) indicate a decrease in R0 and PRCC positive values indicate an increment in R0. Thus, increases in N_v_, N_H_, β_HV_, β_VH,0_, and β_VH,1_ produce a rise in R0 (Fig. 2). The positive effect (PRCC) of these parameters ranges between 0.61 and 0.65 for the model without oral transmission and when it is included, the contribution of vector-host transmission rates (β_VH,0_, β_VH,1_) is shared by both parameters (β_VH,0_ PRCC = −0.30 and β_VH,1_ PRCC = − 0.33). On the other hand, vector and host death rates have significant effect on lowering R0 (model without oral transmission μ_V,_ PRCC = −0.61 and μ_H_ PRCC = − 0.58, including oral transmission μ_V,_ PRCC = −0.61 and μ_H_ PRCC = − 0.58).

**Figure. 2.**
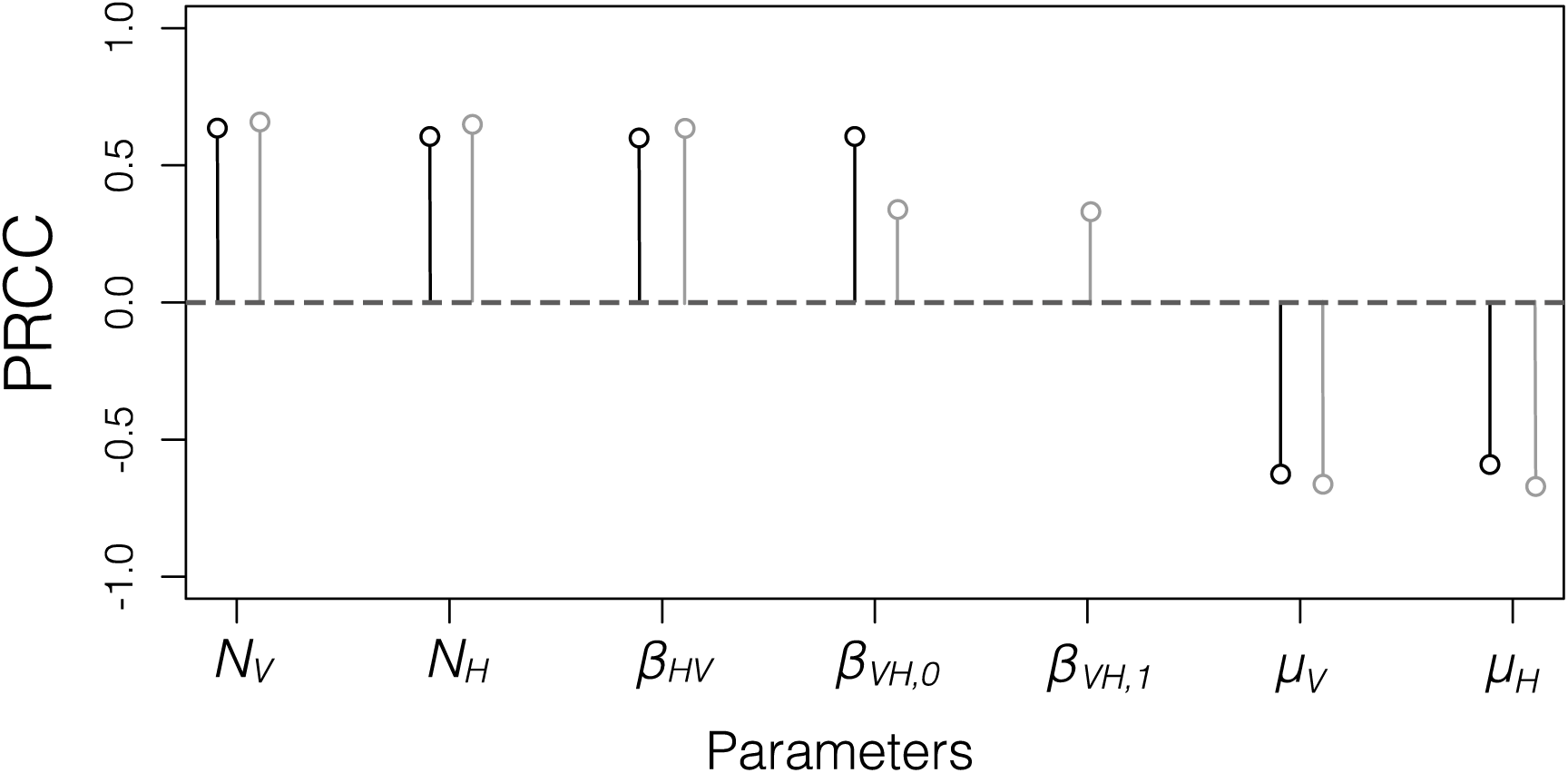
Latin Hypercube Sampling for the global reproductive number R0. PRCC: Partial Rank Correlation Coefficient. Black colored circles correspond to the model without oral transmission and gray to the model including oral transmission. For information about each parameter’s explanation see Table 1. Note that parameters related to host-vector (and vice versa) infection rates and populations (N_v_, N_H_, β_HV_, β_VH,0_, and β_VH,1_) have a positive contribution to R0. On the contrary, vector and host death rates (μ_V_ and μ_H_) have a negative effect in R0.

Plugging the parameter values into the maximum critical virulence equation *μ*(*G*) and considering the maximum probability value in each transmission rate (*β*) we got that *μ*(*G*) max is between 0.3584 and 0.0224. Since none of these values are greater than 0.618, an endemic infection with the parasite cannot occur, interestingly this indicates that another route of transmission, apart from the vector route, is needed to explain high percentages of host infection rates. Figure 3 show a numerical exploration of the model considering parameter ranges for *β*_*HV*_ and *β*_*VH*,0_, here is evident that for every combination of parameters a model that just incorporates vectorial transmission is not capable to exhibit an R0 bigger than one.

**Table I.** Parameters used in the mathematical model of the sylvatic cycle of Chagas disease.

| *Terminology* | *Definition* | *Value* | *References* |
| --- | --- | --- | --- |
| $N_V$ | Total number of vectors<sup>a</sup> | 9.6/palm | [26] |
| $N_H$ | Total number of hosts<sup>a</sup> | 1.5/100ha | [26] |
| $\beta_{HV}^*$ | Max transmission rate of vector by contact with infected host<sup>b</sup> | 1 | [55] |
| $\beta_{VH,0}^*$ | Max transmission rate of host due to an infected vector (biting)<sup>ab</sup> | $1 \times 10^{-2}$ | [45] |
| $\beta_{VH,1}^*$ | Transmission rate of host due to an infected vector (oral)<sup>ab</sup> | $1 \times 10^{-1}$ | [45] |
| $\beta_{HH}^*$ | Transmission rate of host by contact with infected host<sup>b</sup> | $1 \times 10^{-1}$ | [55] |
| $\mu_V$ | Death rate for vectors<sup>a</sup> | 1.7/year | [46] |
| $\mu_H$ | Death rate for host<sup>a</sup> | 0.666/year | [47,48,49] |

\* Transmission rates are reported to the nearest order of magnitude.
<sup>a</sup> Parameter was calculated using literature reports.
<sup>b</sup> Parameter was calculated using expert knowledge.

**Figure 3.**
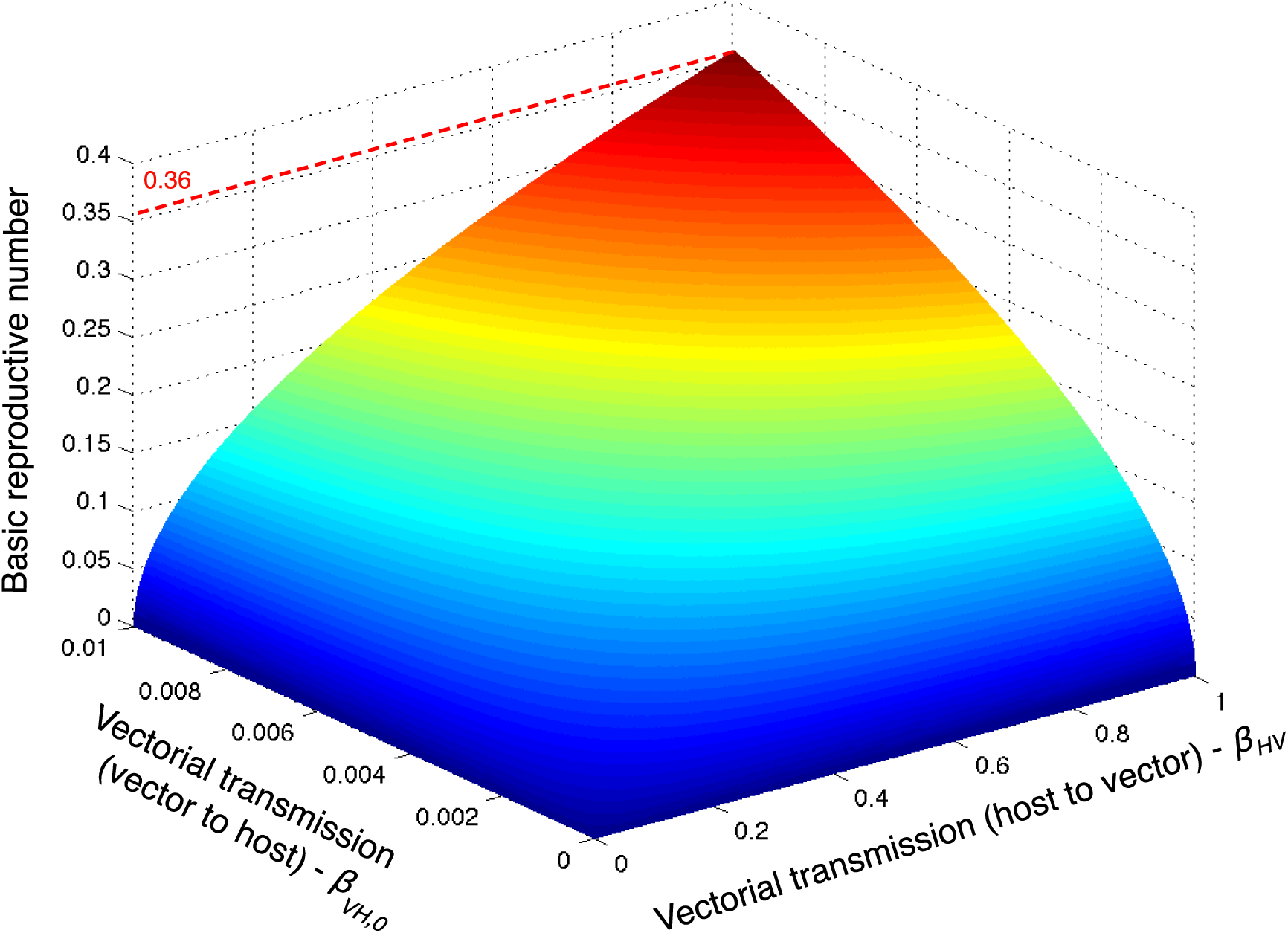
R0 values depicted in z depending on *β*_*HV*_ (here in x) and *β*_*VH*,0_ (here in y) we used for the analyses the palm as the study unit. In this case the max R0 is lower than 0.4.

In addition, Figure 4 shows values of R0 resulted from simulations of the model with a range of values of the oral transmission parameter *β*_*VH*,1_ assuming that *β*_*HV*_ and *β*_*VH*,0_ are at the maximum value found in figure 2, we found that *β*_*VH*,1_ has to be 0.07 or bigger in order to reach and *R*_0_ above 1 (Figure 4).

**Figure 4.**
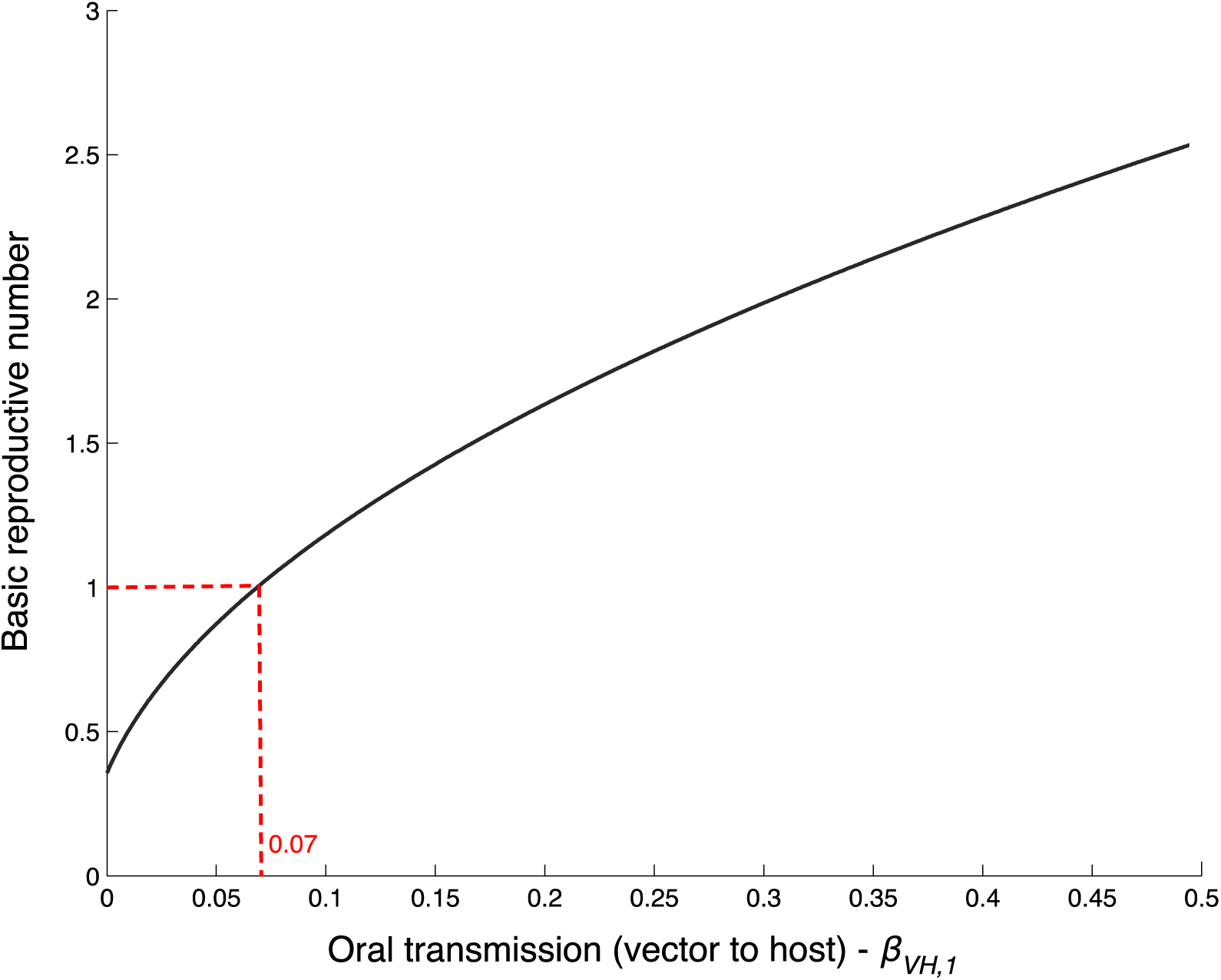
The x axis represents n *β*_*VH*,1_ values range and their corresponding Ro. We assume maximum values for *β*_*HV*_ (1) and *β*_*VH*,0_ (0.01). In this case, *β*_*VH*,1_ must be larger than 0.07, for Ro to have a value above 1.

## Discussion

To the date, studies that investigate in detail the dynamics of the sylvatic cycle of *T. cruzi* transmission are still rare. Nonetheless, it has been shown for the endemic region of Casanare a stable sylvatic transmission, where *R. prolixus* individuals were captured in palm trees (*A. butyracea*) [25-27]. These studies have reported infections rates in mammals ranging from (21-89%) [26, 34, 50, 51]. This is remarkable fact, given that vectorial route has been considered inefficient since the parasite faces great challenges to infiltrate the host bloodstream via vectorial route [52], thus this route its unable to explain the observed reservoir prevalence reported in the literature.

*T. cruzi* vectorial transmission has been suggested to be among one the most inefficient ways for parasites to infect susceptible hosts, although the number of infected hosts it is high on the field fluctuating between 40 to 90% [31-33]. Our results from model simulations only considering vectorial transmission show that the basic reproductive number *R*_0_ is always less than 1. This implies that an additional transmission route is needed to guarantee an endemic state (R0>1). Although, there are no reports in the literature for natural occurring populations where the transmission of the parasite is not supported due to low transmission rates. Perhaps for other vector populations, like *Triatoma dimidiata*, this could be the case and it would be interesting to verify it in the field.

One of our main goals with this study was to establish if vectorial transmission per se was able to explain the high-reported levels of infected hosts. Our results demonstrate that even if we simulate the system at their maximal critical virulence *μ*(*G*) over the highest possible value combinations of transmission rates the system never reaches high number of infected hosts. The incorporation of a new route of transmission, such as the oral transmission, let the system reach the high proportion of infected hosts reported in field studies [47,52-54]. However, these high proportion of infected host could also be obtained with higher levels of transmission rates, in particular increasing the transmission rate from vectors to host *β*_*VH*_, even though we believe that the high values needed are outside of the biological range we have no report to compare to and thus this is mainly speculative.

Results from our sensitivity analysis suggest there is no single variable or parameter that by itself explains the dynamics in the systems. Instead, we were able to identify a subset of factors that together help to explain the temporal variation in the system. First, in the absence of oral transmission the dynamics is explained by the transmission rate and the population size and the incorporation of the oral transmission add a significant positive effect that help to reach a higher number of infected hosts. In addition, it is important to note that the direction of change in the fraction of infected hosts after a change in the parameter is given by the sign of its condition number: a same direction change (e.g. increase parameter-increase *i*_*H*_) is a positive condition number, while an opposite direction change (e.g. increase parameter-decrease *i*_*H*_) produces negative condition numbers. Using the parameter values in table 1, we found that for increasing growth rates increases the fraction of infected hosts. However, the effect of changing *N*_*V*_ is an order of magnitude lower than changing *N*_*H*_ in the same proportion (same relative change). For death rates, we have an opposite-direction effect as it was expected, meaning that if we have more vector or host, the contact between both would be lower accordingly to the mass action law, so the net flow of individuals from susceptible to infected populations would be lower. Again, we saw that vectors have an effect almost an order of magnitude higher than the hosts. Thus, we could expect that these parameters become an interesting target for disease control initiatives.

Importantly, our model is implemented using the palm as spatial unit; the choice is based in its fundamental role in the ecology of both vector and host populations. A mathematical model with a different spatial resolution (i.e. a village) faces the challenge of vector and host mobility. In addition, a temporal model often assumes complete mixing in the spatial component and that is an important aspect to study any host – vector disease model. Here restricting the model to the palm for the analysis simplify and constrain the model results. Extrapolating our results to villages with multiple palms has to be done carefully because hosts often visit multiple palms and insects could move also between palms and houses. We believe that our results should apply to higher levels of aggregation such as villages with high and homogeneous palms density with easy access between palms to guaranteed population mixing. However, including the palm distribution in the model implies a different theoretical approach that although is an important hypothesis it is out of the scope of this paper.

From a biological perspective, the ability of the model to capture important disease dynamics is what makes it useful for testing potential control strategies and studying *cruzi* transmission in sylvatic host species different from *D. marsupialis*. For example, if we increase death rate of vectors and simultaneously decrease the transmission rates to a point where they cancel out is possible to eradicate the disease. This is an important result, because control strategies often target one parameter at a time (i.e. increase vector mortality - house spraying, decrease contact rates - improve house materials), but it seems more reasonable to intervene all of them at the same time taking care in shifting the disease balance from endemic to temporal. In fact, it has been proposed that using a particular control method does not exclude using another one [10]. One way to achieve this at the household level is to combine spraying, which increases insect mortality, with presence of non-reservoirs peri-domestic species, such as chickens, providing another vector feeding. However, there is potential negative effect because the latter could increase vector carrying capacity. In addition to the previous analysis the model could be used to further refine the range of unknown parameters. For example, the transmission rates, *β*_*HV*_ and *β*_*VH*_, are difficult to estimate, however if one has reports of the densities for host and vectors, proportion of infected host and vectors, and a good estimate of death rates, then it is easy to calculate the transmission rates. This technique could be implemented combining fieldwork and the mathematical expressions to make the model adequate to a certain region, and thus a useful disease control tool.

## Conclusions

In Latin America, the transmission dynamics of Chagas disease vary significantly between regions and the Orinoco epidemiological scenario involves a unique mixture of factors that requires interdisciplinary approaches. Computational models, along with biological knowledge, are a great tool to test hypotheses and forecast epidemic events, becoming great allies in understanding transmission mechanisms and designing control strategies.

## Acknowledgements

[funding sources should not be included here or in the manuscript file, only during manuscript submission]

